# Conservation value of low-elevation forests for birds in agroforestry-dominated landscapes in a biodiversity hotspot

**DOI:** 10.1101/2023.08.26.554947

**Authors:** Nayantara Biswas, Siddharth Biniwale, Vishal Sadekar, Yukti Taneja, Kulbhushansingh Suryawanshi, Anand M Osuri, Rohit Naniwadekar

**Affiliations:** Nature Conservation Foundation, 1311, “Amritha”, 12^th^ Main, Vijayanagar First Stage, Mysuru 570017, Karnataka, India; Snow Leopard Trust, 4649 Sunnyside Ave N #325, Seattle, WA 98103, United States

**Keywords:** Functional Diversity, Joint Species Distribution Modelling, Land-use Change, Phylogenetic Diversity, Plantations, Traits, Western Ghats

## Abstract

1. Protected areas (PAs) and multi-use landscapes beyond PAs play important and complementary roles in conserving global biodiversity. Across the tropical forest biome, remnant forests and agroforestry plantations in mixed-use landscapes often have high taxonomic diversity, but their ability to sustain functionally and phylogenetically diverse assemblages and retain conservation-priority species is unclear. Additionally, our understanding of impacts of land-use change across species, function and communities is poor.
2. In India’s Western Ghats (WG) biodiversity hotspot, like in many other tropical regions, PAs cover just 12% of the land and are concentrated in higher elevations. Here, in an approximately 15,000 km^2^ landscape in the northern WG, we compared bird communities across six land-use categories: state-owned high-elevation PAs and low-elevation Reserved Forests beyond PAs, and privately-owned periodically clear-felled low-elevation forests and three types of low-elevation agroforestry plantations (cashew, mango, and rubber).
3. We sampled 184 line transects totalling 137.5 km across habitats, compared taxonomic, functional, and phylogenetic diversity metrics and used joint species distribution models to test for species-trait-habitat associations.
4. Forests in general and lowland forests in particular had higher taxonomic, functional and phylogenetic diversity, and higher occurrence probabilities of evergreen forest-affiliated, conservation-priority, and frugivores than agroforestry plantations. The ranking of land-use types varied across indicators, with rubber ranking higher than cashew and mango for functional and phylogenetic diversity but lower in species occurrence probabilities.
5. Synthesis and Applications: Our findings underscore the irreplaceability of less-disturbed forests for birds in mixed-use landscapes, especially when considering multiple dimensions of biodiversity, habitat specialists, and conservation-priority species. We advocate the simultaneous examination at multiple levels (community, species and traits) to determine the impacts of land-use change as divergent information of conservation importance is gleaned from each level. Reserved and private forests of low elevations of the Northern Western Ghats are critical for avian conservation, given that they harbour significant diversity, and the absence of representative PAs in the region. Given the vulnerability of these low-elevation forests to conversion for non-forestry activities and overexploitation for fuelwood, there is a need to develop viable models of partnership with landowners in these multi-use landscapes for ecological restoration of degraded forests.

## INTRODUCTION

Agricultural expansion is the primary driver of biodiversity loss in the Afro-Asian tropics (Oakley & Bicknell, 2022). Protected Areas (henceforth PAs) are crucial for conserving threatened biodiversity (Cazalis et al., 2020), but have limited geographic coverage (Baldwin & Beazley, 2019). Moreover, PAs are skewed towards higher elevations, particularly in the Afro-Asian tropics (Elsen et al., 2018), leaving low-elevation forests more vulnerable to conversion into agriculture and agroforestry (Namkhan et al., 2021). Therefore, many forest-specialist species typically found in tropical lowlands are more vulnerable to habitat loss than ones at higher elevations (Mills et al., 2023). This underscores the importance of assessing the potential of landscapes beyond PAs – comprising agroforestry plantations and privately- or government-owned forest remnants – for conserving tropical biodiversity.

Biodiversity responses to land-use change have largely been examined through the lens of taxonomic diversity, but this is known to have limitations. For example, shifts in species prevalence and community composition without changes in overall species richness, such as the replacement of specialists by generalist species, might go entirely undetected by taxonomic diversity metrics (Hillebrand et al., 2018). This has led to growing interest in other dimensions of biodiversity, such as functional and phylogenetic diversity (Cadotte et al., 2013). Functional diversity is sensitive to variation in the diversity of functional traits, such as losses of large-bodied animals (Coetzee & Chown, 2016), and phylogenetic diversity to the persistence of evolutionarily-distinct species (Turley & Brudvig, 2016). Moreover, the extent of overdispersion or clustering of communities in functional and phylogenetic space can also shed light on the putative roles of competition and environmental filtering in structuring communities (Cadotte et al., 2013; Webb et al., 2002).

While studies are beginning to examine land-use change impacts on biodiversity through taxonomic, functional, and phylogenetic lenses, much inconsistency exists in the strength and direction of impacts across studies. There are cases where land use change has no effect (Rurangwa et al., 2021), increases (Sreekar et al., 2021), or reduces functional and phylogenetic diversity (Bregman et al., 2016). This highlights a current knowledge uncertainty and the need for more studies examining multiple dimensions of biodiversity from different land-use change contexts and regions. While several studies compare forests with structurally very different land uses, such as farms and urban environments (García-Navas & Thuiller, 2020; Sreekar et al., 2021), there are relatively few studies that contrast the taxonomic, functional, and phylogenetic diversities of forests versus tree-based agroforestry systems (Jayathilake et al., 2021). Addressing this gap is pertinent for Asia, where agroforestry plantations pose pervasive threats to wet tropical forests (Namkhan et al., 2021).

Beyond community-level metrics, assessing species-level responses can provide important insights, especially in the tropics, where many species have narrow geographic ranges, low population densities, and specialised niches (Orme et al., 2006; Stratford & Robinson, 2005). Species’ responses to environmental variables could themselves be influenced by their traits. Examining species-level responses can, therefore, complement community-level biodiversity metrics for understanding the impacts of land-use change and conservation planning. However, studies seldom look at responses at multiple levels of organisation from communities, species and their traits. This is vital for a nuanced understanding of the impact of an anthropogenic driver on community organisation (Leveau et al., 2020).

Birds provide an interesting system to determine the impacts of land-use change on biodiversity. They are good indicators of habitat quality (Canterbury et al., 2000). They perform diverse ecological functions (Sekercioglu, 2006). Their morphological traits are useful for assessing functional diversity as they are strongly related to their niches and ecological function (Pigot et al., 2020). Additionally, land-use and climate change are projected to be responsible for 50% range contractions by 2100 of more than 900 bird species worldwide, and species from the tropics are considered especially vulnerable to land-use change (Jetz et al., 2007).

Here, we assessed taxonomic, functional, and phylogenetic diversity and explored factors structuring bird communities and conservation importance across high-elevation PAs, less-protected lowland forests outside PAs (Reserved and private forests), and lowland agroforestry monocultures (cashew, mango, rubber). Specifically, we asked (1) whether elevation and land-use change filter bird communities, such that lowland, less-disturbed forests have taxonomically more diverse and functionally and phylogenetically less clustered bird communities than high-elevation PA forests, disturbed forests and agroforestry systems; (2) assess species-level responses to elevation and land-use change, with a focus on habitat specialist and conservation-concern species; and (3) which functional traits explain species responses to elevation and land-use change. We report results from the study based on sampling 184 sites in approximately 15,000 km^2^ of the modified landscape in the northern Western Ghats. Previous research on land-use change and birds has focused solely on taxonomic diversity and been concentrated in the southern and central regions of the Western Ghats (Anand et al., 2008; Karanth et al., 2016; Shankar Raman & Sukumar, 2002). By contrast, the northern Western Ghats are an ecologically important and distinctive region that has remained highly understudied (but see Munje & Kumar, 2022). This study presents community-wide, species-specific and trait-specific responses to forest degradation and conversion, thereby providing comprehensive insights for predicting the impacts of one of the most pervasive drivers impacting biodiversity across a diverse range of land uses.

## METHODS

Western Ghats-Sri Lanka Biodiversity Hotspot is one of the ‘hottest’ hotspots due to the high endemism and threats it faces, and it is an area of ‘significant conservation concern’ (Myers et al., 2000). Over 35% of forests in the Western Ghats have been lost in the last 100 years, and forest conversion to agroforestry plantations is one of the primary drivers of forest loss in the Western Ghats (Reddy et al., 2016). We studied in the northern Western Ghats of Maharashtra state, India, between January and May 2022. While the forests in the southern Western Ghats have been mostly converted to tea, coffee and rubber, the privately-owned forests of northern Western Ghats are being rapidly converted to mango, cashew, and rubber (Rege & Lee, 2022). Compared to the southern parts, the northern part of Western Ghats has longer dry spells lasting 7–8 months, making the wet forest species more vulnerable to land-use change. The precipitation and temperature in the region vary between 2150–7450 mm and 11°C–38°C, respectively (Jog, 2009). The terrain is undulating, with the elevation of our sampling sites varying from 12–1110 m a.s.l. The area primarily harboured evergreen and semi-evergreen forests, but they have been degraded or lost due to repeated clearing or conversion to agroforestry plantations (Kulkarni & Mehta, 2013).

We sampled government-owned (PAs and Reserved Forests (henceforth RFs)) and privately-owned forests and cashew, mango and rubber plantations (15.71°–17.76°N, 73.01°–74.71°E; Table 1, Fig. S1). Unlike the privately-owned forests that are clear felled periodically (every 5–10 years) for fuel wood, the RFs and PAs do not experience such drastic and periodic habitat modifications. As per Indian Forest Laws, all forms of resource use are prohibited in PAs, while cattle grazing and fuelwood collection by local communities are permitted in RFs. All the sampling sites in the RFs, privately-owned forests and agroforestry plantations were in lower elevations, while the PA sites were in higher elevations. While the less-disturbed RFs served as reference sites to determine the effects of land-use change on birds, comparing low-elevation RFs with high-elevation PAs allowed us to determine the value of the unprotected low-elevation forests.

**Table 1.** Sampling effort across land-use types. Protected Area sites were in high elevations, while others were in relatively lower elevations. While the three agroforestry plantations are completely modified, the private forests are periodically (5–10 years) clear-felled. The Reserved Forests and Protected Areas are relatively less-disturbed sites.

| Land-use Category | Number of transects | Total sampling effort (km) | Average elevation (± SE) (m) | Raw species richness | Total number of individuals |
| --- | --- | --- | --- | --- | --- |
| Protected Area | 19 | 29.79 | $863 \pm 22$ | 61 | 654 |
| Reserved Forest | 17 | 20.62 | $248 \pm 29$ | 86 | 622 |
| Private Forest | 29 | 22.59 | $346 \pm 53$ | 80 | 734 |
| Cashew | 55 | 23.06 | $153 \pm 7$ | 78 | 814 |
| Mango | 39 | 20.29 | $116 \pm 10$ | 69 | 642 |
| Rubber | 25 | 21.16 | $162 \pm 21$ | 63 | 331 |
| Total | 184 | 137.5 |  | 138 | 3797 |

### Bird sampling

We used line-transect surveys for sampling birds. We sampled 184 sites (one transect per site) across all categories with a total sampling effort of 137.5 km (Table 1). Most plantations were small. Therefore, the transect length varied between 0.17–2.19 km. The total effort in each land-use type was similar (Table 1). Two transects were separated by at least 500 m to minimise spatial autocorrelation. The transects were walked in the morning and afternoon between 0700–1200 hr and 1450–1820 hr, respectively. We recorded bird species identity (sightings and calls) and the number of individuals (when seen). We follow the Clements Checklist of the Birds of the World for taxonomy (Clements, 2007). We only considered terrestrial (forest) bird species seen within 50 m on either side of the transect for the analysis. Since we were interested in determining the impacts of land-use change from forests to agroforestry landscapes on forest birds, wetland species, like egrets, herons and ibises, were ignored from the analysis. We collated functional trait data and phylogenetic data following established methods. Please see Appendix 1 for details.

### Taxonomic, Functional and Phylogenetic Diversity

We used sampling coverage (the probability of not finding a new species with the next sampled individual) to determine sampling completeness (Roswell et al., 2021). In our case, the sampling coverage was high (>90%). To compare species richness across different land uses, we used the Hill-Shannon Diversity measure, standardised for coverage (92% - least for rubber plantations). This diversity measure lays less emphasis on rare species than species richness, is useful for diversity comparisons and is robust to standardisation (Roswell et al., 2021). For sample-coverage-based rarefaction, we used the R-package iNEXT (Hsieh et al., 2016). We bootstrapped the data 50 times to estimate 95% confidence intervals. We interpreted significant differences if the 95% CI did not overlap following Cumming et al. (2007).

We used Mean Pairwise Distance (MPD) to determine functional and phylogenetic diversity. MPD is the average pairwise (phylogenetic/functional) distance between species in a community. With the conversion of forests to agroforestry plantations, we expected the filtering of species to result in a decline in functional and phylogenetic diversity. We used the Standardised Effect Size of MPD since MPD is influenced by species richness. See Appendix 2 for details.

### Species and Trait Responses to Land-use Change

We used the Hierarchical Modeling of Species Communities framework (Ovaskainen & Abrego, 2020) to determine the species’ responses to land-use change and whether functional traits and phylogenetic affinities influence species’ responses. We excluded rare species detected in less than 5% of transects as they provide little information on community assembly and pose challenges for MCMC convergence (Ovaskainen & Abrego, 2020). Thus, we modelled the occurrence of 59 bird species as a function of different land-use categories, the elevation of the transect and trail length. We used the R-package ‘Hmsc’ to fit the model with default prior distributions (Tikhonov et al., 2020). We sampled posterior distribution with three Markov Chain Monte Carlo (MCMC) chains. Each chain was run for 375,000 iterations. We removed the first 125,000 iterations as burn-in. The chains were thinned by 1000 to yield 750 posterior samples for three chains. Please see Appendix 3 for additional details.

We classified the 59 bird species as those preferring evergreen and deciduous forest species based on existing literature (Ali & Ripley, 1999) and those whose populations have remained stable, declined or are uncertain based on their long-term population trends in India as determined by The State of India’s Birds (SoIB) using the eBird data (SoIB, 2020). Long-term population trend data was not available for *Brachypodius priocephalus.* We compared the beta coefficients (from HMSC analysis) across the six land-use categories for the different bird species based on their habitat preferences (evergreen/deciduous) and long-term population trends (declining/uncertain/stable) using the 95% bootstrapped confidence intervals (Cumming et al., 2007).

## RESULTS

### Bird taxonomic, functional and phylogenetic diversity across land-use types

We detected 3797 individuals of 138 bird species across land-use types (Table 1). This included three ‘Vulnerable’ and two ‘Near Threatened’ species. The RFs, which were our reference sites for low elevations, had 1.4–1.8 times higher Hill-Shannon diversity than private forests, the three kinds of agroforestry plantations and the high-elevation PAs (Fig. 1A).

**Figure 1.**
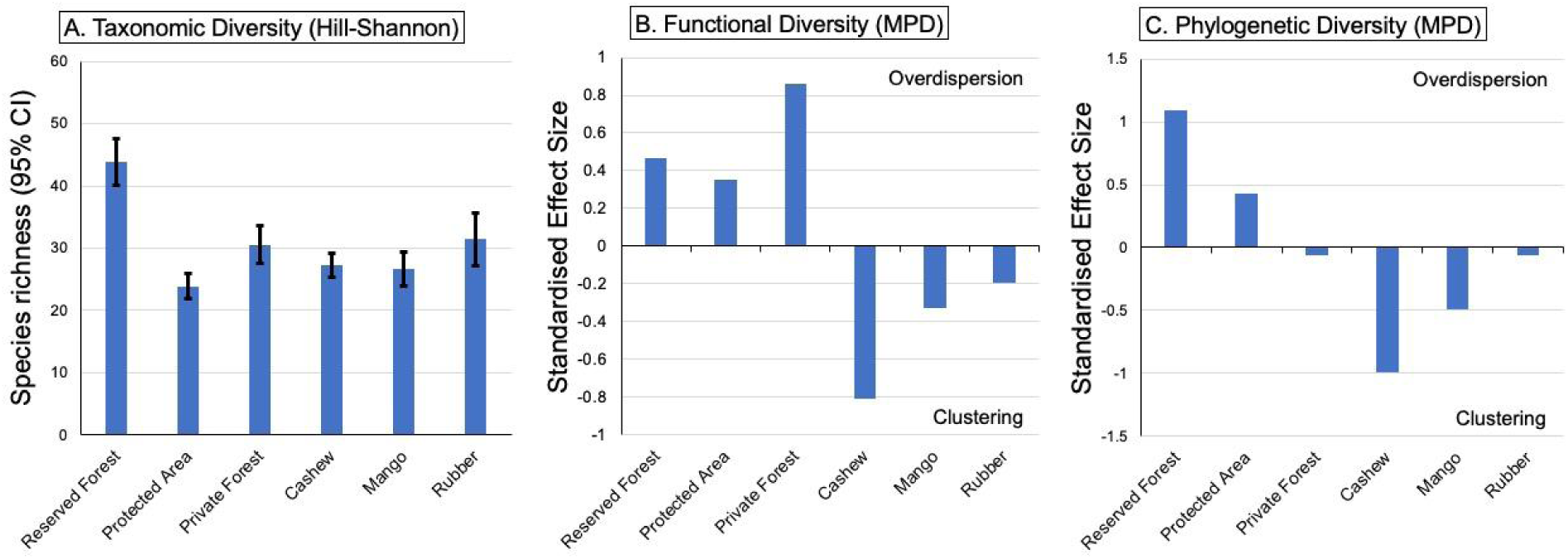
Taxonomic, functional and phylogenetic diversity of birds across the different land-use categories. MPD - Mean pairwise distance. The forested habitats tend towards overdispersion, while the agroforestry plantations tend towards functional and phylogenetic clustering.

The percentage variance explained by the extracted first principal components for trophic, locomotory, diet and strata traits were 98.4%, 91.1%, 62.3% and 54.1%, respectively. The functional and phylogenetic diversity of birds tended to have positive values for less-disturbed forested habitats (PAs and RFs) (Fig. 1B and 1C).

### Species responses to land-use change

The average explained variation in occurrence for all species combined was 20%. For several range-restricted and endemic species found in peninsular India (e.g. *Iole indica*, *Ocyceros griseus*), the total variation explained was greater than 30% (Fig. S2). Among the different fixed effects, land-use type explained more than 50% of the variation in species occurrence, followed by elevation (20.5%) (Fig. S2).

The probability of occurrence of 40 of the 59 species (68%) was lower in at least one of the modified habitats (private forests or agroforestry plantations) than in the reference RF sites (Fig. 2A). The coefficients of overall bird species responses for agroforestry plantations was lower than forested habitats as inferred from non-overlapping 95% bootstrapped CIs (Fig. 2B) indicating negative impacts of agroforestry plantations on birds. Compared to low-elevation RFs, the probability of occurrence of 20 species (e.g. *Ocyceros griseus*, *Dicrurus aeneus*) was lower in all modified habitats (Fig. 2). Compared to RFs, the probability of occurrence of 10 species (e.g. *Leptocoma minima*, *Spilornis cheela*) was lower in all the three types of agroforestry plantations. Some of these species (e.g. *O. griseus*, *B. priocephalus*, *M. horsfieldii*, and *L. minima*) are range-restricted (found only in peninsular India) or Western Ghats endemics. The occurrence of several open habitat species (e.g. *Pycnonotus cafer*, *Cinnyris asiaticus*) was higher in agroforestry plantations than in the reference RF sites (Fig. 2). Probability of occurrence of only nine species (11%) (e.g. *Columba elphinstonii*, *Hypsipetes ganeesa*) was higher in high-elevation PAs than the low-elevation RFs (Fig. 2).

**Figure 2.**
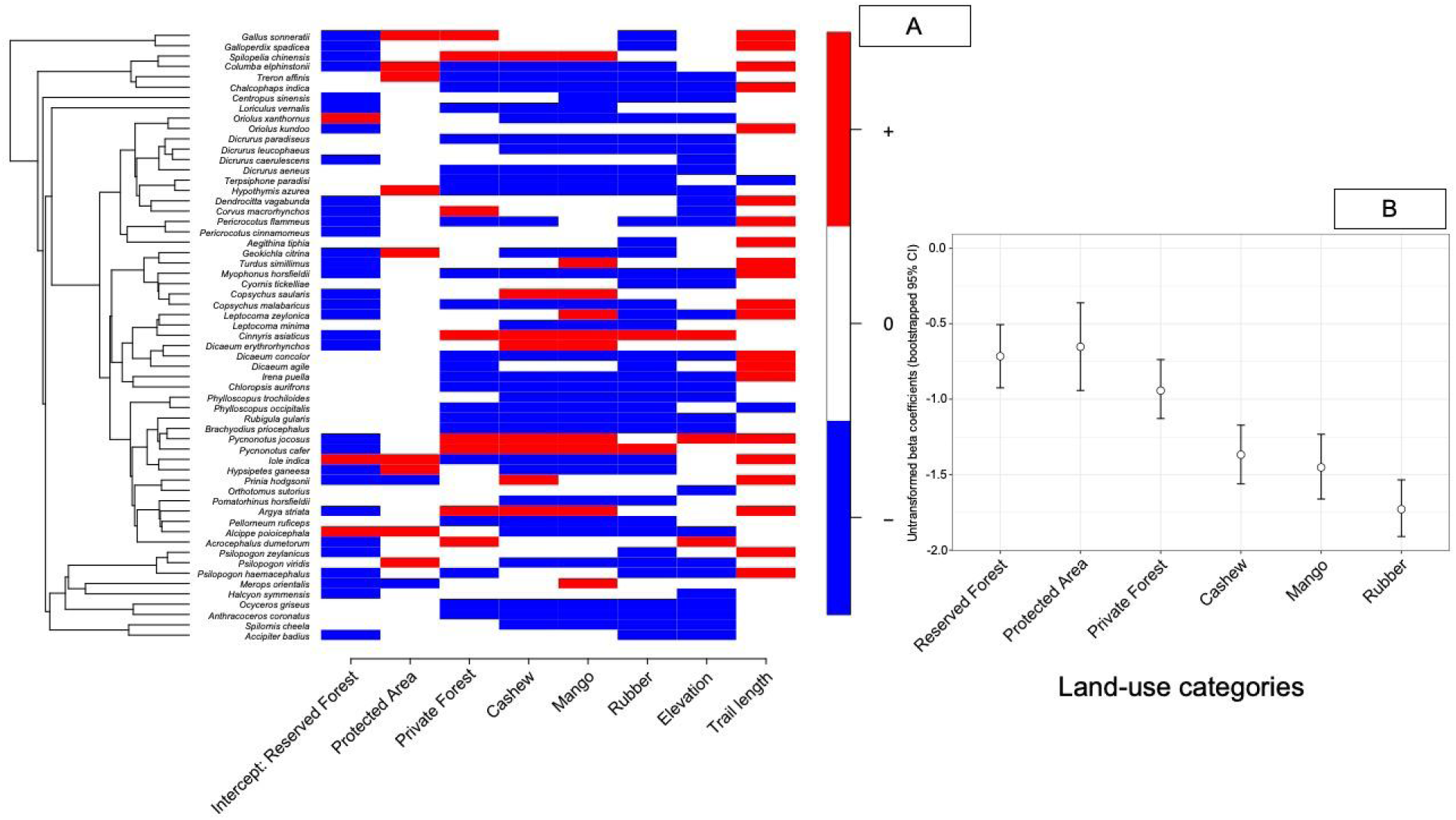
A) Species-specific responses of the 59 bird species to different predictors. Blue and red indicate negative and positive associations with posterior probability ≥ 0.95, respectively. The probability of occurrence of many species was lower in modified landscapes than in the Reserved Forests. B) Comparison of mean beta-coefficients (untransformed) and associated bootstrapped 95% CI for birds across different land-use types. The estimated means were the least for rubber.

### Responses of evergreen forest specialists and species with declining populations

The probability of occurrence of species that preferred evergreen forests was lower in the agroforestry plantations than in forested habitats as inferred from non-overlapping bootstrapped 95% CI (Fig. 3A). There was no discernible pattern for species that preferred the open deciduous forests (Fig. 3B). The probability of occurrence of species showing declining trends as per the State of India’s birds was lower in agroforestry plantations (Fig. 3C). The pattern was not evident for species having stable or uncertain population trends (Fig. 3D and 3E). Among agroforestry plantations, the probability of occurrence of birds with stable or uncertain trends was lower in rubber plantations than the benchmark low-elevation RFs (Fig. 3D and 3E).

**Figure 3.**
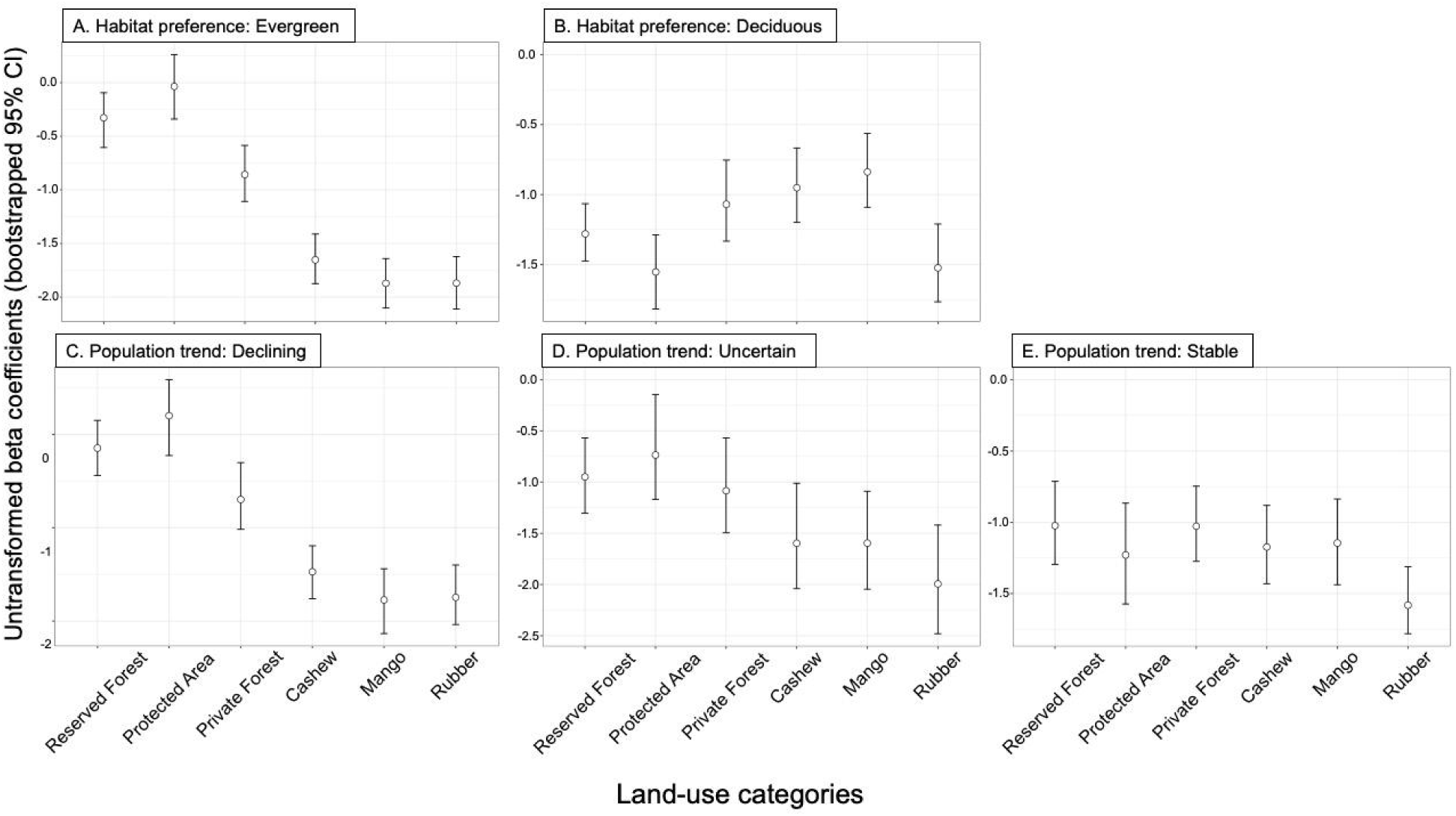
Untransformed beta coefficients and bootstrapped 95% CI for each land-use type as obtained from Hierarchical Modelling of Species Communities for birds classified into different groups. A-B: coefficients in different land-use types for birds found in evergreen forests and those not found in evergreen forests. The mean coefficient in modified landscapes (including Private Forests) for evergreen species is lower than in forested ecosystems. C-E: mean coefficients in different land-use types for birds that have exhibited declines in population trends over time (C), whose population trends are uncertain (D), and whose populations have remained stable as determined by the State of India’s Birds (2019). Populations that have exhibited long-term declines in populations in India are negatively affected by habitat modification.

### Traits and species response to land-use change

Traits explained 20% of the variation in bird species responses to different predictor variables. Frugivorous diet was negatively associated with land-use change (Fig. 4). High strata values were associated with birds that forage on the ground or lower strata (e.g. *Galloperdix spadicea*, *Copsychus saularis*) and low with canopy-dwelling birds (e.g. hornbills). Strata was negatively associated with the reference RFs, indicating a greater prevalence of canopy-dwelling birds in RFs (Fig. 4). We did not find any evidence of a phylogenetic signal in species-response to predictor variables (⍴ = 0; 95% CI: 0.0-0.4), indicating that closely-related species did not respond more similarly than unrelated species to predictors beyond what was expected from their functional traits.

**Figure 4.**
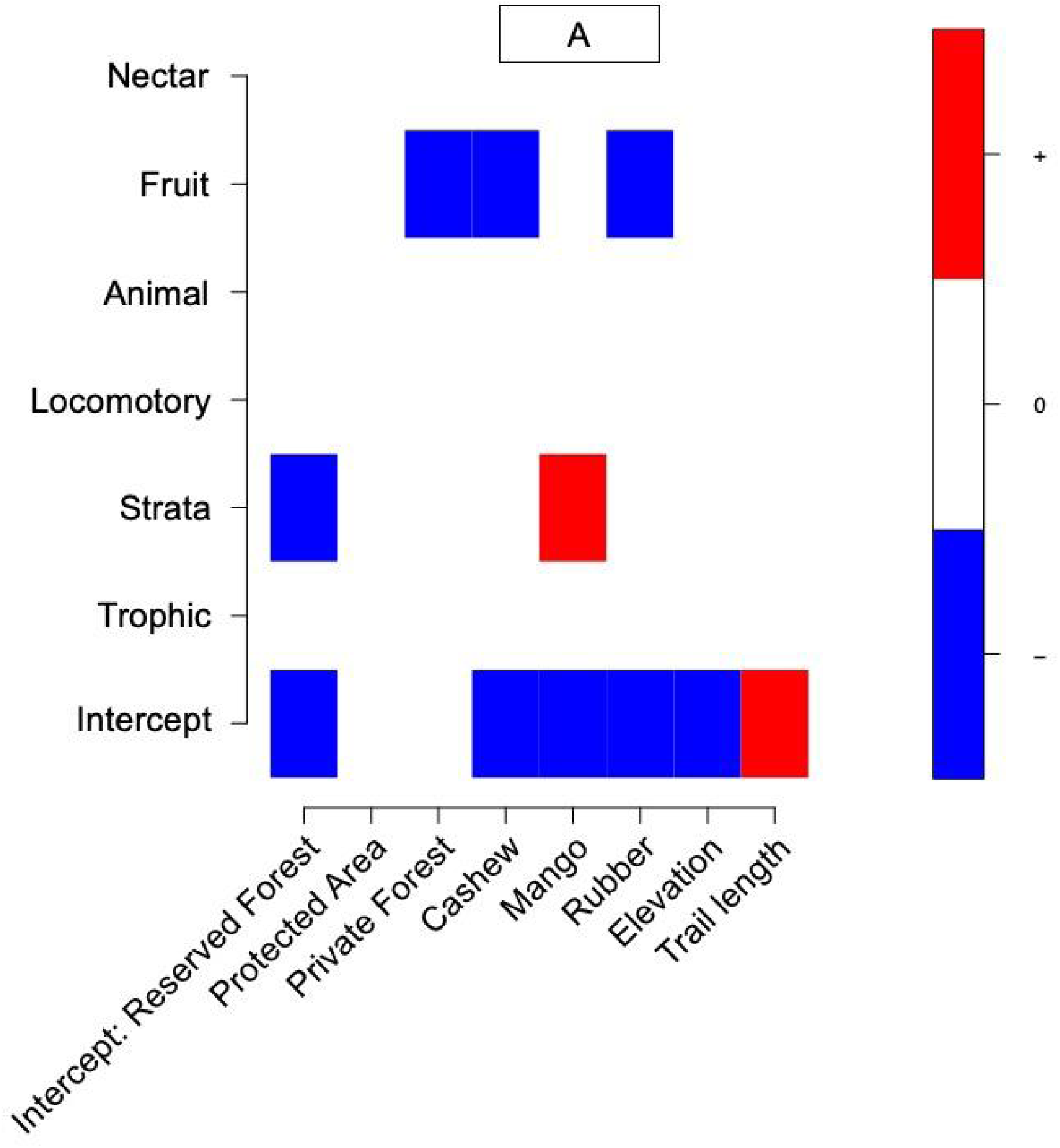
Mean posterior gamma parameter values that determine the influence of species traits on species response to different predictor variables. Blue and red indicate negative and positive associations with posterior probability ≥ 0.95, respectively.

## DISCUSSION

We examined community-, trait- and species-level responses of birds to land-use change by comparing three forested ecosystems (varying in protection level and elevation) and three agroforestry plantations using spatially replicated transect surveys in the Western Ghats Biodiversity Hotspot. We found evidence for the decline of several evergreen forest species and frugivores in privately-owned forests that are periodically disturbed and agroforestry plantations. In some cases, the loss of evergreen forest species was countered by deciduous forest species. However, it was still insufficient to maintain diversity, as modified landscapes tended towards clustering while the two less-disturbed forested ecosystems (PA and RFs) tended towards overdispersion. The low-elevation RFs had a higher diversity of birds than the high-elevation PAs, highlighting the value of securing remnant forest patches in lower elevations for biodiversity conservation. Collectively, our results highlight the importance of forests beyond PAs for lowland tropical avifauna, and suggest the need for collaborative conservation and restoration models to secure these forests.

Previous studies have mostly focused on determining land-use change impacts on taxonomic, functional and phylogenetic diversity (Chapman et al., 2018; Penjor et al., 2022) and, on occasion, functional traits (Sreekar et al., 2021). In this study, along with community metrics and functional traits, we have also determined individual species responses and responses of forest specialists and species with large-scale population declines to land-use change. The value of integrating data at multiple levels of organisation (community, traits and species) is particularly evident when we closely examine the impacts of rubber plantations. At the community level, rubber might appear more bird-friendly than other agroforestry plantations, but this is not supported at the species level, where species occurrence probabilities declined more in rubber plantations. The number of detections of birds was far lower in rubber plantations than in other plantations and forested habitats. Thus, not all agroforestry plantation types are similar, and some, like rubber plantations, may be more detrimental for birds in this region. This is corroborated by another study that has demonstrated a lower density of birds across foraging and habitat guilds in rubber plantations than coffee and areca (Karanth et al., 2016). Unlike the mango and cashew plantations, rubber plantations are relatively more recent in the landscape (Munje & Kumar, 2022), and legacy effects could potentially boost their functional and phylogenetic diversity (García-Navas & Thuiller, 2020; Hendershot et al., 2020). Just investigating community-level metrics would not have likely revealed the finer details of the impacts of rubber plantations on birds. We recommend that future research should focus on simultaneously examining species-, trait- and community-level responses to land-use change to ensure that negative impacts of land-use change are not masked because of examination at a single level.

Contrary to some of the previous studies that have not consistently found lower functional and phylogenetic diversity of forest birds in agroforestry landscapes than in forests (Penjor et al., 2022; Rurangwa et al., 2021; Sreekar et al., 2021), we find that modified landscapes had lower taxonomic diversity, and functionally and phylogenetically clustered communities. Conversion of forests to open farmland landscapes results in the loss of forest birds and the addition of wetland birds (Prescott et al., 2016; Rurangwa et al., 2021; Sreekar et al., 2021). The addition of wetland birds may compensate for the reduction in functional diversity because of the loss of forest birds. This likely results in inconsistent patterns in functional diversity with drastic land-use change, as observed in earlier studies. We focused on forest birds and compared tree-crop landscapes (and not open agricultural landscapes) with degraded and less-disturbed forests. Therefore, we may have found reduced functional and phylogenetic diversity in modified landscapes.

Conversion to modified landscapes resulted in losing functionally important groups, like frugivores. The occurrence of hornbills, wood-pigeon, fairy bluebird and multiple endemic bulbuls, all of which play important roles in seed dispersal (Naniwadekar et al., 2019), was negatively associated with degraded private forests and agroforestry plantations (Fig S3). Large frugivores (e.g. hornbills and *Ducula* pigeons) play functionally unique roles. They are critical for the seed dispersal of large-seeded plants in the Asian tropics (Naniwadekar et al., 2021; Sethi & Howe, 2009). These species are critical for forest regeneration, and their loss can alter the recovery of degraded forests in the landscape.

Agricultural landscapes and monocultures typically have lower phylogenetic diversity than forests, suggesting phylogenetic filtering and loss of evolutionarily distinct lineages (Morelli et al., 2018). Reduction in phylogenetic diversity is driven by the loss of species and increased relatedness between species (Frishkoff et al., 2014). The occurrence of members of several distinct lineages (e.g. hornbills, bluebird, leafbird, leaf warblers) was lower in altered landscapes (Fig. S3). In other cases, the loss of distinct genera within a lineage (e.g. *Rubegula*, *Iole*, *Bradypodius* among bulbuls) in altered landscapes were associated with the replacement by fewer closely-related species (e.g. *Pycononotus* spp.). Thus, the observed reduction in phylogenetic diversity and associated phylogenetic clustering of the community resulted from the loss of evolutionarily distinct lineages and increased relatedness between the species that persist in altered landscapes.

Broadly, bird diversity declines with increasing elevation with functionally and phylogenetically clustered communities in higher elevations on tall mountains (Montaño-Centellas et al., 2020; Quintero & Jetz, 2018). Higher elevations associated with low temperatures may impose physiological and energetic barriers to persistence (Barve et al., 2021; Jankowski et al., 2013). While some species appear to respond negatively to elevation gradients, and we detected a reduction in species richness, the cumulative effect did not translate into a signal of phylogenetic or functional clustering of the communities. However, the 1100 m elevation gradient we examined in the northern Western Ghats is narrower than the Himalayan or Andean gradient, which could explain the lack of phylogenetic clustering with increasing elevation in our study.

Most PAs in the northern Western Ghats are restricted to higher elevations with very poor representation in the lower elevations. We demonstrate that the low-elevation forests harbour higher diversity. They also harbour significant populations of threatened species (Mudappa & Raman, 2009; Pawar & Sadekar, 2023). The mean expected probability of occurrence of Malabar Grey Hornbill (Vulnerable), Malabar Pied Hornbill (Near Threatened) and Grey-headed Bulbul (Near Threatened) was higher in the low-elevation RFs than in the high-elevation PAs. This highlights the importance of remnant patches of RFs. Unfortunately, the private forests are degraded due to periodical (5-10 years) clearing (Biswas et al. in prep.). Ratnagiri is one of the two districts we sampled that harbour low-elevation forests. Unlike the Sindhudurg district, where 10% of the district’s geographical area is under RFs, only 1.3% of more than 60% of the forested land in Ratnagiri district is under the RFs (Kulkarni & Mehta, 2013). Most forests here are privately owned. The government-owned RFs are vulnerable to diversion for non-forestry use and infrastructure development (LIFE, 2020). Given the importance of the RFs in the area for harbouring avian biodiversity and the vulnerability of private forests to land-use change, the existing RFs should be declared as Conservation Reserves and their denotification should be avoided. Declaration of some of the RFs as Conservation Reserves has already been achieved in the region, thereby setting a precedent for remaining patches of RFs (https://indianexpress.com/article/cities/mumbai/maharashtra-tillari-area-in-sindhudurg-declared-conservation-reserve-6473276/).

### Conservation Implications

Given the differences in outcomes of community, species and trait level responses, studies should investigate land-use change impacts on biodiversity at multiple levels. This also provides information on responses of evolutionarily distinct and conservation concern species to land-use change, which can inform management decisions. While conversion to agroforestry plantations may result in similar structural changes to forests and may negatively impact endemic species and those of conservation concern, different agroforestry plantations differ in their impacts on communities and species. Studies often examine biodiversity responses to different forms of agriculture in consolidation by pooling information across land-use categories (farms, plantations). However, community and species responses must be examined for each kind of agricultural practice separately. This is important as different agricultural practices may differ in their management regimes (e.g. undergrowth and leaf litter management and use of pesticides and weedicides) with varying impacts on biodiversity. More than 100 km^2^ of forests are diverted for non-forestry use annually in India, most of which is in low elevations. Linear infrastructure and mining or quarrying are responsible for a significant proportion of forest diversion (LIFE, 2020). Relatively less-protected RFs are more vulnerable to forest conversion than PAs. Given that most of the land is privately owned in the study region, the remnant patches of RFs are vital reservoirs of biodiversity. Suitable alternatives to the diversion of RFs are vital in this region. These low-elevation forests in Asia are critical habitats for threatened birds. There need to be systematic efforts to identify remnant low-elevation forests and accord them better protection. Collaboration between landowners and conservation practitioners is vital for protecting and restoring degraded forests on private lands (Sarnaik et al., 2017).

## AUTHOR CONTRIBUTIONS

RN, AMO and KS conceived the ideas and designed the methodology; SB, NB, and VS collected the data; RN, NB and YT analysed the data; RN, NB, YT and AMO led the writing of the manuscript. All authors contributed critically to the drafts and gave final approval for publication. Our study brings together authors from different states within India, where the work was conducted. Four of the seven authors (SB, VS, KS and RN) are from the state (Maharashtra) where the study was conducted. Whenever relevant, literature published by scientists from the region has been cited.

## ACKNOWLEDGEMENTS

We are grateful to the Maharashtra Forest Department, particularly Sunil Limaye (CWLW), Nanasaheb Ladkat, Uttam Sawant, Vishal Mali, DFOs of Sawantwadi and Chiplun Forest Division, Suhas Patil and Ajit Mali for giving us the necessary permissions (No: Desk-22(8)/WL/Research/CR-53(20-21)/3361/22-22) and support to conduct fieldwork. We thank On the Edge Conservation (UK) for funding this work. We thank Ninad Gosavi, Himanshu Lad and Siddhi Damle for helping with fieldwork. We are grateful to Suri Venkatachalam, Madhura Niphadkar, Atul Joshi, Aparna Watve, Praveen Desai, Milind Patil, Parag Rangnekar, Hemant Ogale, Vishwas (Bhau) Katdare, Girish Punjabi, Shashank Dalvi, Kaka Bhise, Prasad Gavde, Mahesh Mhangore, Gajanan Shetye, Vinayak Sapkal, Pooja Pawar and Anushka Rege for support and discussions. We thank all the villagers and households who warmly hosted us during our travels.

## DATA ACCESSIBILITY STATEMENT

The data will be uploaded on Dryad (www.datadryad.org) on acceptance of the manuscript.

## Notes

### Competing Interest Statement

The authors have declared no competing interest.

