## Appendix 1; Fig. S* for "Conservation value of low-elevation forests for birds in agroforestry-dominated landscapes in a biodiversity hotspot"

**SUPPLEMENTARY MATERIAL**

**APPENDIX 1**

**Functional and Phylogenetic Data**

To estimate overall functional diversity in each land-use type, we focused on combining morphological and ecological traits following Chapman et al., (2018). Morphological traits included trophic and locomotory traits, and ecological traits included their diet and foraging strata. The morphological traits were sourced from AVONET bird trait database (Tobias et al., 2022). The ecological traits were sourced from EltonTraits 1.0 (Wilman et al., 2014). We used beak length (culmen), width, and depth to capture trophic traits, and tarsus length, wing length, Kipp's distance and tail length to capture locomotory traits (Chapman et al., 2018). We used diet and foraging strata traits from "EltonTraits 1.0" (Wilman et al., 2014) following (2020). The multiple ecological traits are represented as percentages in columns for every species. The diet categories used are percentages of invertebrates, vertebrates, fish, carrion, fruits, nectar, seeds and other plant materials in the diet. The categories for foraging strata include information on the prevalence of foraging on the ground, in the understory, mid-high, in the tree canopy and aerial (Wilman et al., 2014). To reduce the dimensionality of the data, we performed the Principal Component Analysis, following Chapman et al., (2018) and (2020). The extracted first principal components (PCs) for each of the four trait categories (locomotory, trophic, diet and strata) were then scaled to make species x traits matrices for each land-use category. To create a functional dendrogram of traits, we used R packages "vegan" (Oksanen et al., 2019) and "ade4" (Dray & Dufour, 2007).

To determine the effect of land-use change on the phylogenetic diversity of birds, we selected a phylogenetic tree with maximum phylogenetic support from a distribution of 100 randomly generated trees (Ericson backbone) downloaded from global bird phylogeny (www.birdtree.org) (Jetz et al., 2012).

### APPENDIX 2

#### Functional and Phylogenetic Diversity Analysis

Standardised effect size (SES) calculates (Observed mean - Expected mean)/ SD (expected), where SD is the standard deviation of the randomised assemblage. Negative SES values (observed value lower than the null mean) are indicative of clustering, and positive values (observed value greater than the null mean) indicate overdispersion (Swenson, 2014). We calculated SES using an independent swap algorithm (Gotelli, 2000), where the probability of drawing a species from the assemblage depends on its overall abundance in the data. We used the "picante" package (Kembel et al., 2010) in R to calculate SES MPD. We expected the general trend in SES values to show over-dispersion (due to competition) in forests and under-dispersion (due to environmental filtering) in plantations

### APPENDIX 3

#### HMSC analysis

We included the length of the trails as a fixed effect in the model since the lengths of transects were variable. We included the location of each trail as a random effect to account for potential sample non-independence. Since habitat modification, especially conversion to agroforestry plantations, can alter the vegetation structure and resource availability for birds, we expected traits linked to stratum, locomotion, trophic level and diet to be impacted by land-use change. We obtained information on the percentage contribution of animal matter, fruits and nectar in the diet of different bird species from Elton Traits 1.0 (Wilman et al., 2014). We used beak length (from culmen), beak width and beak depth traits to capture trophic functionality, tarsus length, wing length, Kipp's distance and tail length traits to capture locomotory functionality, and percentage foraging on ground, understory, mid canopy, tree canopy and above vegetation to capture strata-specific functionality in birds following Chapman et al., (2018) and Montaño‐Centellas et al., (2020). The dimensionality of the multiple data for each trait category was reduced with Principal Component analysis following  Chapman et al., (2018). The extracted principal components (PCs) were then scaled and used in the analysis. To determine the phylogenetic signal in the residual variation of species responses to land-use change, we used a phylogenetic tree with maximum phylogenetic support from 100 randomly-selected trees (Ericson backbone) that were downloaded from global bird phylogeny (www.birdtree.org) (Jetz et al., 2012).

We assessed model convergence through potential scale reduction factor and effective sample size for all the beta and gamma coefficients. We used Tjur $R^2$ to assess model fit (Tjur, 2009) and variance partitioning approach to assess the proportion of variation explained by the three types of fixed effects (land-use type, elevation and trail length), apart from the random effect. We evaluated species' responses to land-use change by examining beta coefficients and the influence of traits on species responses to predictor variables by examining gamma coefficients. We estimated the variation in species responses to the different predictor variables that the traits could explain. We used the rho estimate to determine the phylogenetic signal in species responses. We considered coefficients' influence significant if the 95% credible intervals on beta, gamma and rho parameters did not overlap zero.

### APPENDIX 4

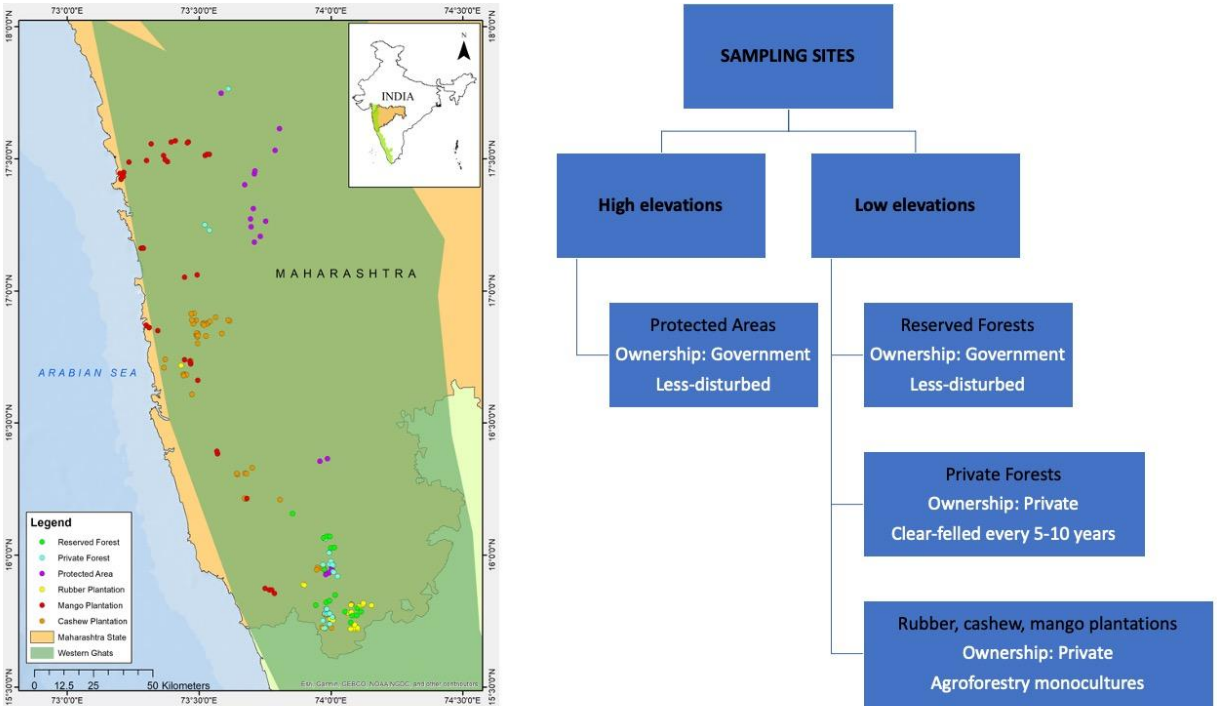


**Figure S1.** Map of the study area. The minimum distance between sampling locations was 500 m. The rubber plantations are mostly in the southern part of the sampling area. There are fewer cashew plantations in the northern portions of the sampling area, where mostly mango is planted.

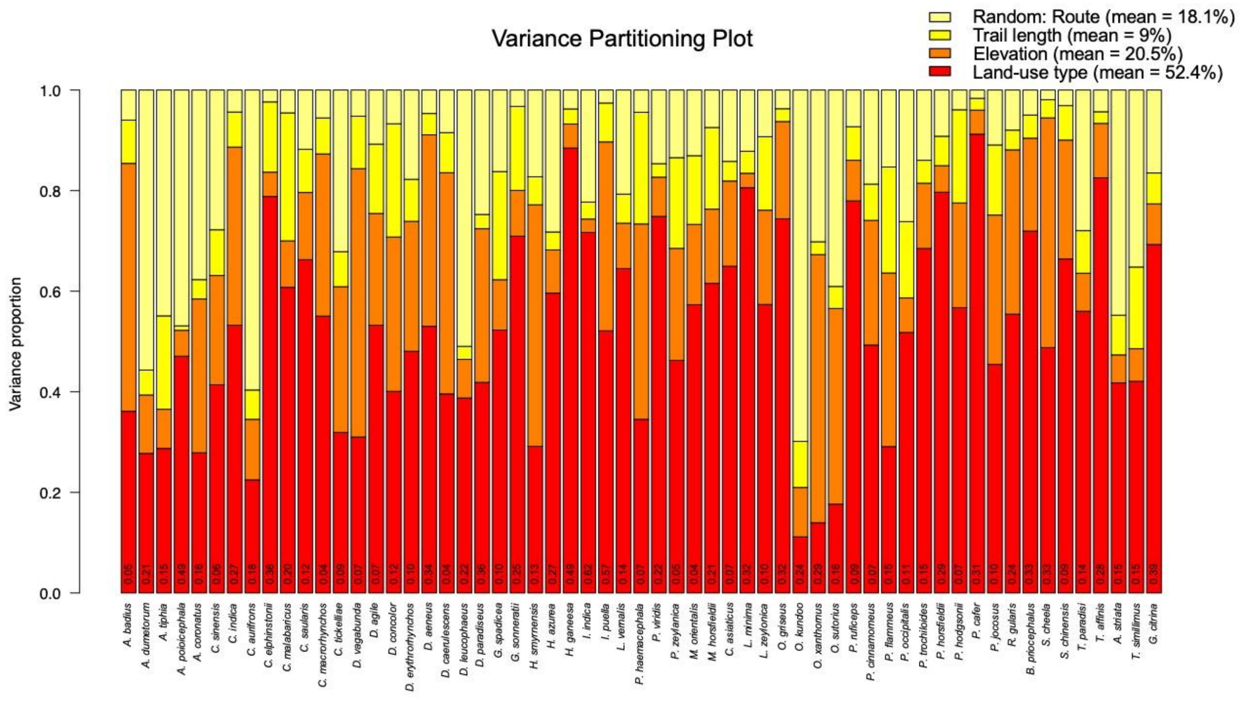


**Figure S2.** Variance partitioning plot for each of the 59 species. Percentage variation explained by the different fixed (land-use type, elevation and trail length) and spatial random effects are summarised in the top right. The Tjur $R^2$ for each bird species is reported in each bar.

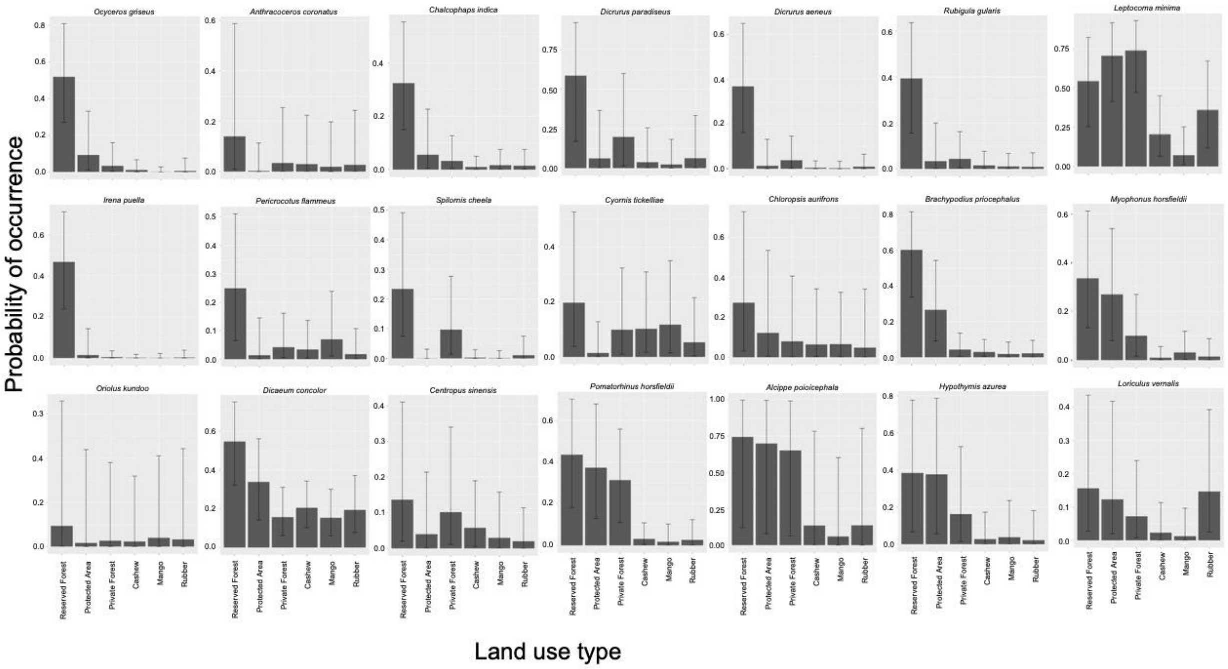
Probability of occurrence
Ocyceros griseus
Anthracoceros coronatus
Chalcophaps indica
Dicrurus paradiseus
Dicrurus aeneus
Rubigula gularis
Leptocoma minima
Irena puella
Pericrocotus flammeus
Spilornis cheela
Cyornis tickelliae
Chloropsis aurifrons
Brachypodius priocephalus
Myophonus horsfieldii
Oriolus kundoo
Dicaeum concolor
Centropus sinensis
Pomatorhinus horsfieldii
Alcippe poioicephala
Hypothymis azurea
Loriculus vernalis
Reserved Forest
Protected Area
Private Forest
Cashew
Mango
Rubber
Land use type

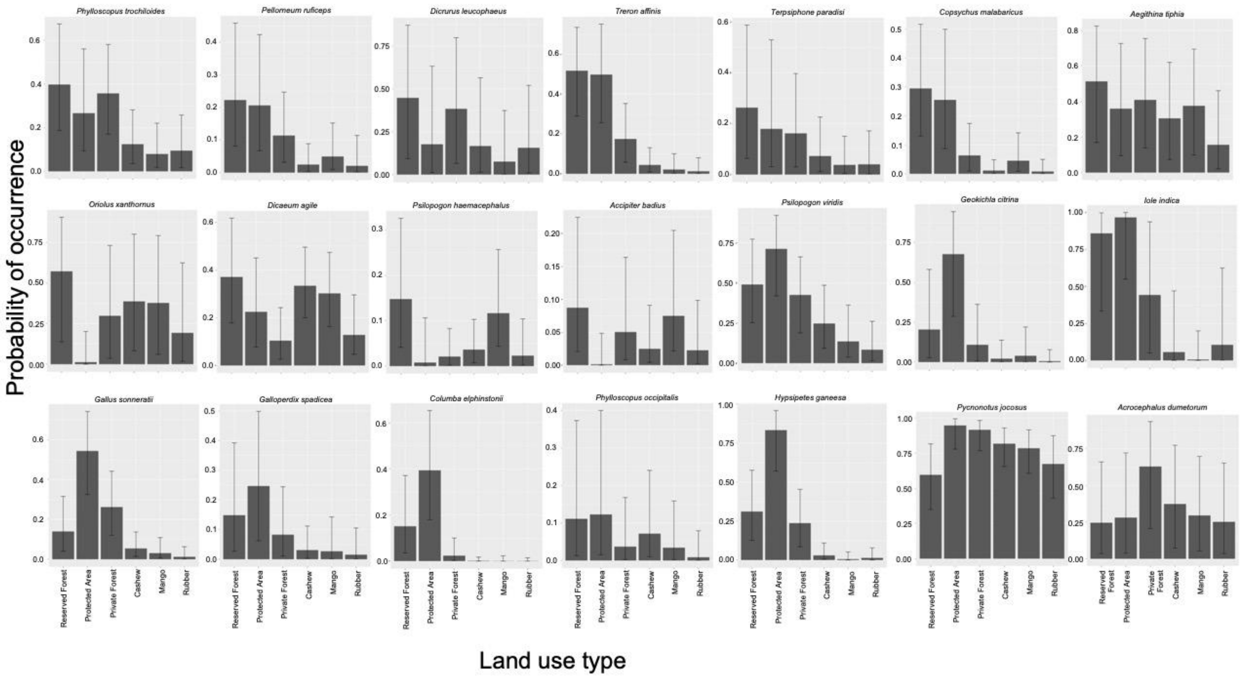
Probability of occurrence
Phylloscopus trochiloides
Pellorneum ruficeps
Dicrurus leucophaeus
Treron affinis
Terpsiphone paradisi
Copsychus malabaricus
Aegithina tiphia
Oriolus xanthornus
Dicaeum agile
Psilopogon haemacephalus
Accipiter badius
Psilopogon viridis
Geokichla citrina
Iole indica
Gallus sonneratii
Galloperdix spadicea
Columba elphinstonii
Phylloscopus occipitalis
Hypsipetes ganeesa
Pycnonotus jocosus
Acrocephalus dumetorum
Reserved Forest
Protected Area
Private Forest
Cashew
Mango
Rubber
Land use type

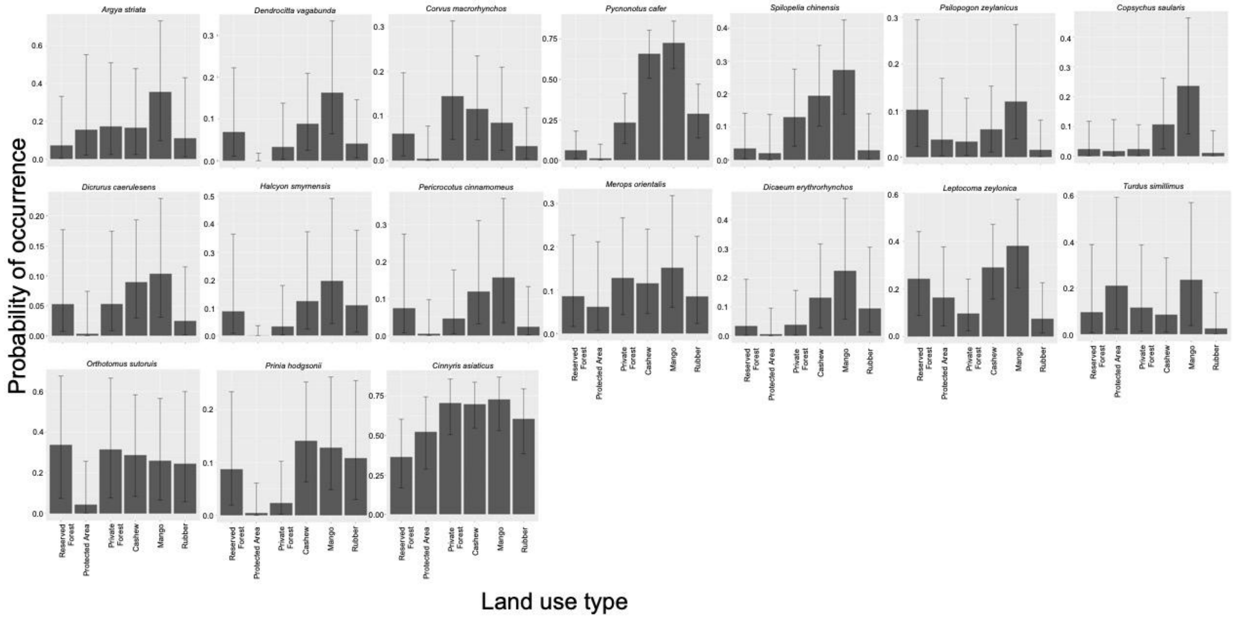


**Figure S3.** The panels show the predicted probability of occurrence across different land-use types for the 59 bird species. Panels A and B are species whose mean probability of occurrence is greater in forested habitats than in agroforestry plantations. Panel C are mostly species whose mean probability of occurrence is greater in agroforestry plantations than in forested habitats.
